# Novel start codons introduce novel coding sequences in the human genomes

**DOI:** 10.1101/2020.11.04.368936

**Authors:** He Zhang, Yang Xie

**Affiliations:** Quantitative Biomedical Research Center, Department of Population and Data Sciences, University of Texas Southwestern Medical Center, Dallas, TX 75390

## Abstract

Start-gain mutations can introduce novel start codons and generate novel coding sequences that may affect the function of genes. In this study, we systematically investigated the novel start codons that were either polymorphic or fixed in the human genomes. 829 polymorphic start-gain SNVs were identified in the human populations, and the novel start codons introduced by these SNVs have significantly higher activity in translation initiation. Some of these start-gain SNVs were reported to be associated with phenotypes and diseases in previous studies. By comparative genomic analysis, we found 26 human-specific start codons that were fixed after the divergence between the human and chimpanzee, and high-level translation initiation activity was observed on them. The negative selection signal was detected in the novel coding sequences introduced by these human-specific start codons, indicating the important function of these novel coding sequences. This study reveals start-gain mutations are keeping appearing in the human genomes during the evolution and may be important sources altering the function of genes which may further affect the phenotypes or cause diseases.

## Introduction

Genetic mutations play key roles in the evolution of genes via providing resources for novel functions. Mutations can cause genetic polymorphism in the population, and contribute to the genetic diversity of the individuals (Auton et al. 2015). During the evolution, some mutations were fixed in the species and became species-specific genetic markers that contribute to both genetic and phenotypic differences distinguishing different species (Varki and Altheide 2005). Single-Nucleotide Variant (SNV) is the most common type of variation in the population, and more than 80 million SNVs have been genotyped in large-scale genetic studies (Auton et al. 2015). Many SNVs can alter the function of the gene by changing the protein sequence and are associated with phenotype diversities and diseases (Buniello et al. 2019). Several kinds of protein-altering SNVs were reported frequently in genetic studies, including missense SNVs (Pal and Moult 2015), nonsense SNVs(Yngvadottir et al. 2009), read-through SNVs (Shibata et al. 2015), and splicing-relevant SNVs (Kurmangaliyev et al. 2013). Besides these SNVs, start-gain SNVs (Cingolani et al. 2012), which was located in the 5’ untranslated region (5’UTR) of the mRNA, can also change the protein sequences via converting triple nucleotides to a novel start codon before the original start codons of the coding sequence (CDS).

The translation of coding sequence in mRNA to protein is a key step in the central dogma, and the start codon plays an important role in translation initiation (Clark and Marcker 1966). The stat-gain SNV can introduce a novel CDS before the original CDS, which may alter the function of this gene, and some start-gain SNVs are associated with human diseases (Semler et al. 2012; von Bohlen et al. 2017). However, the start-gain SNVs were not studied as commonly as other types of protein-altering SNVs, because such SNVs were usually annotated as non-coding SNVs in 5’UTR. During the evolution, a start-gain SNV can be fixed in the population and became a species-specific start codon. However, there was no systematic genome-wide study on the human species-specific start codons.

In this study, we tried to investigate the novel start codons in the human genomes at both population-level and species-level. First, we wanted to know how many potential start-gain SNVs can be found in the natural human populations, and whether they were active in translation initiation. Second, we wanted to know whether there were human-specific start codons generated and fixed in the human genome after the divergence with the chimpanzee.

## Result

### Start-gain SNVs exist in the natural human populations

From 62 Yoruba individuals, 110 potential start-gain SNVs were identified, each possessed 13 to 26 novel start codons in the genome (Supplementary Table 1). For each start-gain SNV, the 62 Yoruba individuals can be split into two groups according to whether the individual had a start-gain allele or not. Then we examined the ribosome occupancy for the sites with start-gain allele and without the start-gain allele. To facilitate the analysis, only 86 start-gain SNVs that passing the following three criteria were used in the ribosome occupancy analysis: 1) at least five codons between the NSC and original start codon, 2) at least three individuals possessed NSC allele and 3) from autosomes. Based on the aggregated results across 86 start-gain SNVs, the ribosome occupancy on the novel start codons in the individuals with the start-gain allele was four times higher than the homologous sites in the individuals without the start-gain allele (P = 0.0093) (Figure 1). For the sites upstream of the novel start codons, the ribosome occupancy was extremely low, which means almost no ribosome was stacked before the novel start codons. In contrast, the sites following the novel start codons showed some signal of ribosome occupancy but not as much as the novel start codons. That was a common pattern observed around the canonical start codons. Instead, the ribosome occupancy around the sites without NSS alleles didn’t show a similar pattern (Figure 1), and more uniformed ribosome occupancy was observed across different positions, which may represent random background noises. This observation suggests some novel start codon indeed have the potential to accumulate ribosomes and further initiate the translation.

**Figure 1.**
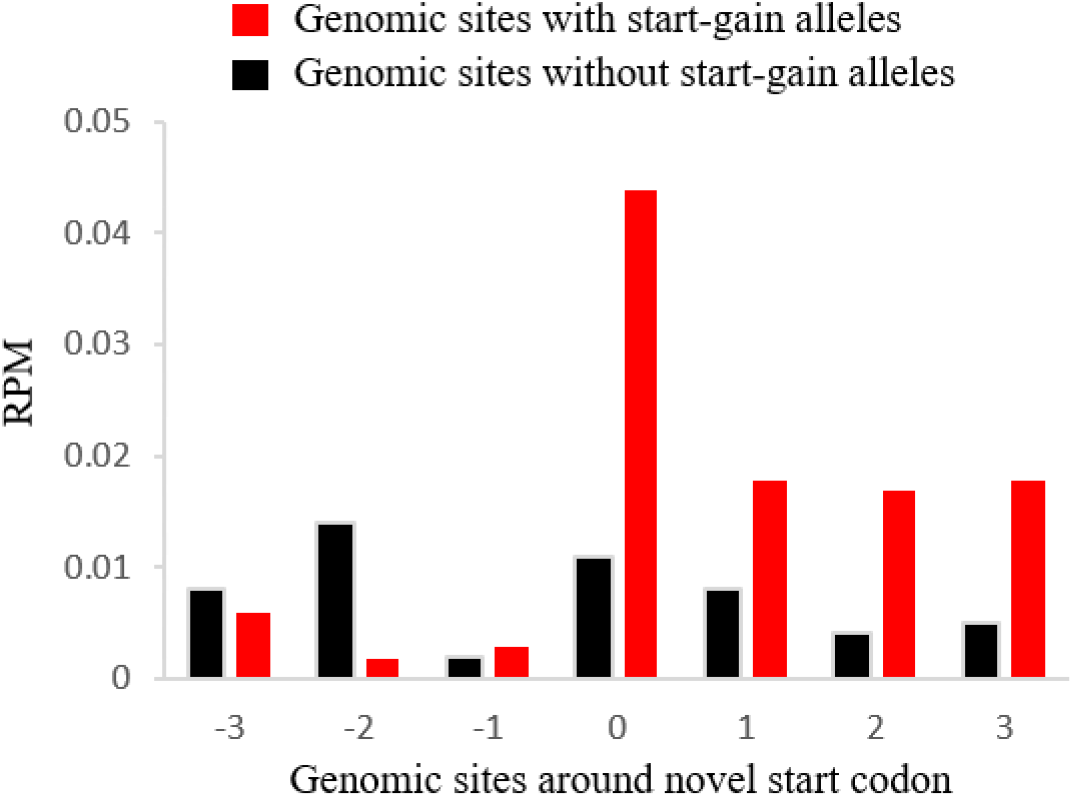
Ribosome occupancy around the start-gain SNVs in the 62 Yoruba individuals. For each polymorphic start-gain SNV, some individuals (red) possessed the start-gain allele introducing a novel start codon and the other individuals (black) possessed an allele not introducing a start codon. Each position represents a codon or a segment of triple-nucleotides. ‘0’ represents the novel start codons (AUG) for individuals with start-gain allele or the corresponding triple-nucleotides for the individuals without the start-gain allele. Y-axis represents the mean RPM of ribosome occupancy aggregated across all start-gain SNVs

Then we investigated the distribution of novel start codons in the natural populations using the dataset from the 1000 Genomes Project. Across all 2,504 individuals from 26 populations, 829 SNVs can introduce a potential novel start codon, and the frequencies of the start-gain alleles were from 0.019% to 99.94% (Supplementary Table 2). Although around 78.5% of these start-codon SNVs were rare (allele frequency less than 0.1%) in the populations, there were still 69 start-gain SNVs with a frequency greater than 1% and 32 of them were common (>5%) in the populations. On average, each individual carried 23 candidate novel start codons in the genome, and that number was 15 if we only considered the common start-gain SNVs with allele frequency greater than 5% (Supplementary Table 3). The median length of the novel coding sequences between the novel start codons and the original canonical start codons was 48nt (16 codons), and some of them can even reach hundreds of codons (Supplementary Table 2).

Interestingly, 12 start-gain SNVs were reported associated with diseases or phenotypes according to the records in the ClinVar database (Landrum et al. 2018), but all of these SNVs were annotated as non-coding SNVs in 5’UTR (Supplementary Table 4). Based on the finding in this study, a potential explanation can be provided for the effect of these SNVs since these potential start-gain SNVs may introduce novel start codons and novel peptides which may affect the function of the protein products.

### Novel start codons were generated and fixed in the human genome after the divergence between the human and chimpanzee

Start-gain SNPs are polymorphic in the human populations, and can only be found in some individuals. Next, we wanted to investigate whether some novel start codon mutations have been fixed in the human populations. By a comparative genomics analysis of the human, chimpanzee, gorilla, and orangutan, 26 human-specific start codons were identified in the human genome (Supplementary Table 5). These start codons were fixed in the human populations but not observed in the chimpanzee, gorilla, and orangutan genomes, which means they were fixed in the human genome after the divergence between human and chimpanzee. The length of the novel coding sequence between the human-specific start codon (hSSC) and the ancestral start codons (hASC) varied from 3nt to 186nt, with a median value of 30nt (10 codons). To test whether the human-specific start codons initiate the translation, ribosome occupancy was evaluated for each human-specific start codon and their orthologous sites in the chimpanzee genome. 16 of 26 human-specific start codons had at least five codons between the human-specific start codon (hSSC) and the ancestral start codon (hASC), and they were used in the ribosome occupancy analysis.

For the ribosome-profiling dataset from 62 Yoruba individuals, the aggregated ribosome occupancy at the human-specific start codons (hSSC) was more than 4.9 times higher than the ancestral start codons (hASC) which were the orthologous sites of the start codons of the chimpanzee genome (Wilcoxon signed-rank test, *P* = 7.4×10^−16^) (Figure 2b, 2c). A similar pattern was observed for four Yoruba individuals from the other independent ribosome-profiling dataset (P = 0.0043) (Supplementary Figure 1a, 1b), confirming the pattern was not due to the random effect in different experiments. That suggests more ribosomes were accumulated at the human-specific start codons (hSSC) than the ancestral start codons (hASC) in some genes, and the human-specific start codons (hSSC) were more active in the translation initiation than the ancestral start codons (hASC). Then we examined the ribosome occupancy in the chimpanzee genomes, and the ribosome occupancy at the orthologous sites of human-specific start codons (cOSC) was not higher than the start codons (cSC) (Figure 2d, 2e). Combining the pattern of ribosome occupancy for both the human and chimpanzee, we can conclude that the orthologous sites of the human-specific start codons in chimpanzee (cOSC) were not as active as human-specific start codons (hSSC) in recruiting ribosomes to initiate the translation.

**Figure 2.**
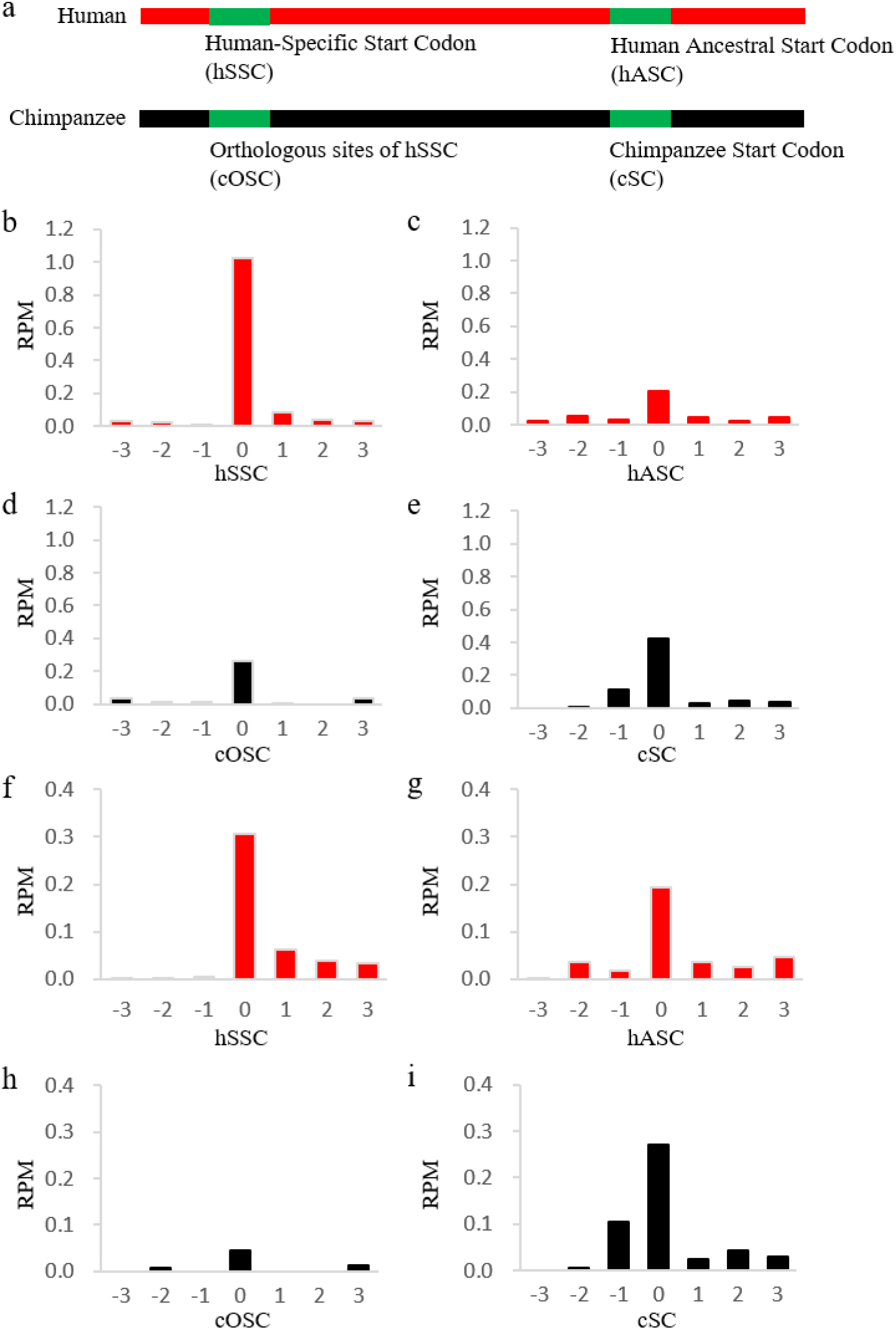
Ribosome occupancy for the human-specific start codons and adjacent positions in the 62 Yoruba individuals and 5 chimpanzee individuals. a) The diagram shows the sites evaluated for the ribosome occupancy. The mean ribosome occupancy was calculated across b) 16 human-specific start codons (hSSC), c) the corresponding downstream human ancestral start codons (hASC), d) the orthologous sites in the chimpanzee (cOSC), and e) start codons in the chimpanzee (cSC). After excluding human-specific start codons originated from CUG, the mean ribosome occupancy was calculated across f) 14 non-CUG originated human-specific start codons (hSSC), g) the corresponding downstream human ancestral start codons (hASC), h) the orthologous sites in the chimpanzee (cOSC) and i) start codons in the chimpanzee.

In the chimpanzee genome, the ribosome occupancy at the orthologous sites (cOSC) of human-specific start codons still was higher than the nearby region, although it was lower than the canonical AUG start codons (cSC). We found that the ribosome occupancy at cOSC sites was majorly contributed by the CUG codon, which was known as the non-AUG start codon in mammalian genomes (Starck et al. 2012). Considering that situation, we excluded two novel AUG start codons converted from CUG in the next round analysis of ribosome occupancy. For the remaining 14 human-specific start codons, the ribosome occupancy in the human-specific start codons (hSSC) was still higher with the ancestral start codons (hASC) (Wilcoxon signed-rank test, *P* = 1.3×10^−5^), but the ribosome occupancy at the orthologous sites of the human-specific start codons in chimpanzee (cOSC) was much lower than that on the canonical start codons (cSC) (Wilcoxon signed-rank test, *P* = 2.4×10^−4^) (Figure 2h, 2i; Supplementary Figure 1c, 1d). That suggests the non-CUG orthologous sites of human-specific start codons (cOSC) have very low activity of translation initiation, while the human-specific start codons (hSSC) showed a strong activity comparable with ancestral start codons (hASC). These observations supported the hypothesis that some human-specific start codons acquired significant activity to initiate the translation after mutated from their orthologous sites in the chimpanzee genome.

### Negative selection acts on the new coding sequences introduced by the human-specific start codons

Since we observed significant translation initiation activity in the human-specific start codons, the novel peptides may be generated by the novel coding sequences between the human-specific start codons (not included) and the ancestral start codons (not included). If these novel peptides were generated in the translation elongation and have a function in the protein product, the evolution constraints can be expected on these novel peptides. To test that hypothesis, we examined the negative selection signals on these novel CDSs.

For each novel CDS following the human-specific start codons, the alignment with its orthologous sequences in chimpanzee and gorilla was extracted from the genome alignment. Since the number of mutations observed in each CDS was limited, alignments for 15 CDSs, which is longer than 15nt, were combined to make a more reliable estimation of the mutation rates among braches. Although the orthologous sequences of these novel CDSs probably are not coding sequences in chimpanzee and gorilla, we still treated them as ‘coding sequence’ to inspect the possible selection during the evolution.

Branch models from PAML (Yang 2007) were used to test whether the negative selection signals can be observed on the CDSs introduced by the human-specific start codons. In the first model, a fixed Ka/Ks value (ω_0_) was assumed across all branches (Figure 3a) to represent the null hypothesis that the Ka/Ks value was the same for each branch. In the second model, a foreground Ka/Ks value (ω_1_) was assigned to the human branch, and the other branches shared the same background Ka/Ks value (ω_0_) (Figure 3b). In the second model, the estimated ω_1_ was 0.36 for the human branch (Figure 3c), while the ω_0_ was 2.74 for other branches (Figure 3d). The Chi-squared test (*P* = 0.028) showed that the second model with two Ka/Ks values is significantly better than the first model with a fixed Ka/Ks ratio (ω_0_=0.97) across all branches. That indicates purifying selection has been operating in these human-specific CDSs to exclude nonsynonymous mutations altering the protein sequence. To check whether we can observe a similar pattern in the chimpanzee and gorilla branches, we also tested the third model in which the chimpanzee branch was assigned a foreground ω_1_ and the fourth model in which the gorilla branch was assigned a foreground ω_1_. In both models, the ω_1_ was greater than one (Supplementary Figure 2c and 2d), indicating no overall negative selection signal was observed on the chimpanzee or gorilla branch. That suggests some of the peptides generated from the CDSs introduced by human-specific start codons may play important functions in the final protein product, and the nonsynonymous mutations that change the protein sequences were not favored during the human evolution. That provides another layer of evidence that human-specific start codons functional in the translation initiation, and the novel CDSs following the novel start codons are functional.

**Figure 3.**
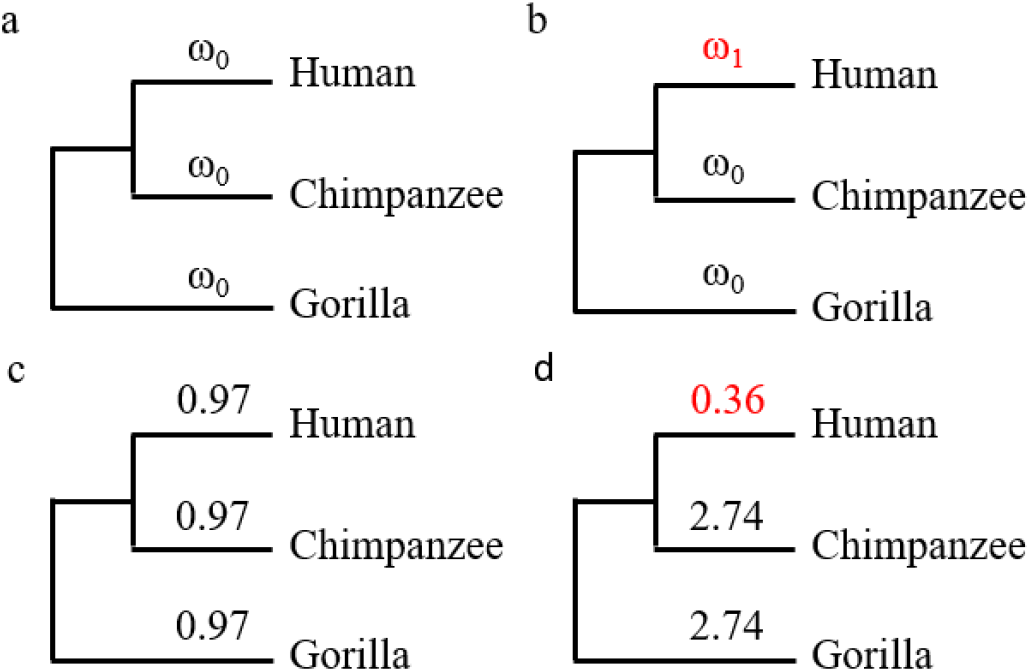
Different branch models were compared to address whether the human branch had a lower ω value than the other branches. a) The first model assumed a fixed ω value (ω_0_) across all branches. b) the second model assumes the human branch had an independent ω value (ω_1_) and the other branches shared the same ω value (ω_0_). The values of ω were estimated for c) the first model and d) the second model using PAML.

## Discussion

In this study, we investigated the novel start codon in species level and population level. Novel start codons were generated continuously during evolution. 26 novel start codons have been fixed in the human genomes after the divergence with the chimpanzee and became a part of genetic mutations distinguishing between the human and other species. Besides that, we observed 829 polymorphic SNVs that potentially introduced novel start-codons in genomes of different modern human populations. These SNVs were not fixed in the human genomes and contributed to the genetic diversity of human populations. Ribosome-profiling results strongly indicate that some of the novel start-codon SNVs in human populations and the fixed human-specific start-codons were active in translation initiation.

Mutations are important sources to generate novel phenotypes or functions in evolution. Novel start-codon SNV can change the sequence of the protein product directly by adding an extra peptide to the N-terminal of the original protein sequence, which may alter the function of the gene directly. Interestingly, several start-gain SNVs were reported to be associated with phenotypes or diseases, and these SNVs were annotated as non-coding mutations in 5’UTR. By assuming that these SNVs may introduce a novel start codon and an extra peptide that changed or disrupted the function of these genes, we got a candidate interpretation of the genetic mechanism for the association between these SNVs and phenotypes. That’s another layer of the evidence that the start-gain SNVs affected the function of the genes.

It’s helpful to detect potential evolution selection signal to infer the potential role of these novel peptides generated by the coding sequences following the novel start codons. Although we cannot assay the evolution signal for each novel coding sequence due to the limited mutations observed after the divergence between the human and chimpanzee, we still observed the negative selection signal if the novel coding sequences from different human-specific start codons were combined. That suggests at least some of the novel coding sequences play an important role in corresponding genes.

Although non-AUG start codons were discovered in the mammalian genomes, we still only consider AUG as the start codon in this study since the non-AUG start codons are rare in the mammalian genomes. We did found that the CUG start codon had considerable activity in translation initiation of some genes, but the other potential non-AUG start codons showed extremely low-level activity. Even in the situation where the AUG was converted from CUG, the translation ignition activity on AUG was still much higher than the CUG as a start codon. Thus it’s reasonable to consider AUG converted from other triple nucleotides as a candidate novel start codon in most situations. If a non-AUG triple became to AUG, that doesn’t necessarily mean it became an active start codon, because usually some other sequence features such as Kozak sequence (Kozak 1987) may be required to promote the translation initiation. However, it’s not clear what features are essential to initiate the translation. Thus we didn’t require the existence of any of those features to define a novel start codon. That would improve the sensitivity to identify the candidate novel start codons, although some of the false positive ‘novel start codons’ may be identified mistakenly.

## Methods

### Identification of candidate novel start codon SNVs in the human populations

All of the genome positions in this study were based on the human reference genome GRCh38 (Schneider et al. 2017). Genotypes were get from the 1000 Genomes Project (Auton et al. 2015). The annotation of the human genome was from GENCODE v30 (Frankish et al. 2019). Firstly, we identified all SNVs that changed non-AUG triple nucleotides to AUG in the 5’UTR of a transcript. Then an SNV can be kept only if the AUG was in the same reading frame with the downstream coding sequence and no stop codon was observed between the AUG and downstream original start codon. Furthermore, SNVs located in any known coding sequences were excluded. The remaining SNVs were considered as the start-gain SNVs.

### Identification of human-specific start codons

Pairwise genome alignments (Syntenic Net) between the human (hg38) and chimpanzee (panTro6), gorilla (gorGor4), and orangutan (ponAbe3) were downloaded from the USCS genome browser (Lee et al. 2020). The annotation of the human genome was from GENCODE v30 (Frankish et al. 2019), and the annotations for the genomes of chimpanzee (NCBI.105) were from NCBI RefSeq (O’Leary et al. 2016).

To reduce the possible false positives caused by misalignment and non-orthologous alignment, only the ‘one-to-one’ alignments (reciprocal single hit) were kept. Based on the filtered genome alignments, we identified all mutations which have different alleles between the human and chimpanzee genomes. If a mutation was located in the annotated start codon of a transcript and its homologous sequence in the chimpanzee genome was within the 5’UTR of a transcript, it was identified as a candidate mutation that introduced a human-specific start codon. Then the alleles at the mutation position in the human genomes were compared with its orthologous alleles in gorilla and orangutan, and a mutation was dropped if either gorilla or orangutan had the same allele as the human. Then we examined the remaining mutations in 2,504 individuals from the 1000 Genomes Project, and only the ones that were monomorphic in all of the 2,504 individuals were considered as fixed in the human populations. Finally, only the mutations fixed in the human populations were kept, and the start codons generated by those remaining mutations were considered as the human-specific start codons.

### Processing of ribosome profiling dataset

The ribosome profiling data of 62 Yoruba individuals were from dataset GSE61742 (Battle et al. 2015) of the NCBI GEO database. The ribosome profiling data of four Yoruba individuals and five chimpanzee samples were from dataset GSE71808 (Wang et al. 2018) of the NCBI GEO database.

All reads were mapped to the curated non-coding RNA (including rRNAs, tRNAs, and snRNAs) (Quast et al. 2013; Chan and Lowe 2016; Kuksa et al. 2019) databases using STAR (2.7.1a) (Dobin et al. 2013), and the reads mapped to any non-coding RNA were dropped. Then, the remaining reads were mapped to the reference genome using STAR, and only the uniquely mapped reads with a length between 26nt and 32nt were kept. The starting position of P-site for each read was estimated using the 12th nucleotide from 5’ end for the reads with a length between 26nt and 29nt and the 13th nucleotide from 5’end for the reads with a length between 30nt and 32nt. Read counts at P-sites were normalized to the unit ‘reads per million’ (RPM) across the genome to represent the ribosome occupancy at every single nucleotide. Obvious periodicity was observed for the normalized reads count at coding sequences (Supplementary Figure 3). To calculate the ribosome occupancy for each codon, the RPMs at the first nucleotide of the codon, the second nucleotide of the codon, and the nucleotide before the first nucleotide of the codon were added up to tolerate error in the cleavage of ribosome footprint.

### Estimation of evolution rate

PAML (4.9j) (Yang 2007) was used to estimate the ratio between nonsynonymous mutation rate and synonymous mutation rate, which is known as Ka/Ks or ω. Different branch models were compared to address whether the human branch had a higher Ka/Ks value than the other branches. The first model assumed a fixed Ka/Ks value (ω_0_) across all branches (Figure 3a), and the second model assumes the human branch had an independent Ka/Ks value (ω_1_) and the other branches shared the same Ka/Ks value (ω_0_) (Figure 3b). The third and fourth models assumed the chimpanzee branch or the gorilla brand had an independent Ka/Ks values correspondingly (Supplementary Figure 2a and 2b). The significance of the difference between the likelihoods of models was evaluated using the Chi-squared test.

## Acknowledgments

This work is supported by the National Institutes of Health [P30CA142543, R35GM136375, 1R01GM115473], and the Cancer Prevention and Research Institute of Texas [RP180805].

## Disclosure Declaration

The authors declare no competing interests.

## Notes

### Competing Interest Statement

The authors have declared no competing interest.

## Reference

Auton A, Brooks LD, Durbin RM, Garrison EP, Kang HM, Korbel JO, Marchini JL, McCarthy S, McVean GA, Abecasis GR. 2015. A global reference for human genetic variation. Nature 526: 68–74.

Battle A, Khan Z, Wang SH, Mitrano A, Ford MJ, Pritchard JK, Gilad Y. 2015. Genomic variation. Impact of regulatory variation from RNA to protein. Science 347: 664–667.

Buniello A, MacArthur JAL, Cerezo M, Harris LW, Hayhurst J, Malangone C, McMahon A, Morales J, Mountjoy E, Sollis E et al. 2019. The NHGRI-EBI GWAS Catalog of published genome-wide association studies, targeted arrays and summary statistics 2019. Nucleic Acids Res 47: D1005–D1012.

Chan PP, Lowe TM. 2016. GtRNAdb 2.0: an expanded database of transfer RNA genes identified in complete and draft genomes. Nucleic Acids Res 44: D184–189.

Cingolani P, Platts A, Wang le L, Coon M, Nguyen T, Wang L, Land SJ, Lu X, Ruden DM. 2012. A program for annotating and predicting the effects of single nucleotide polymorphisms, SnpEff: SNPs in the genome of Drosophila melanogaster strain w1118; iso-2; iso-3. Fly (Austin) 6: 80–92.

Clark BF, Marcker KA. 1966. The role of N-formyl-methionyl-sRNA in protein biosynthesis. J Mol Biol 17: 394–406.

Dobin A, Davis CA, Schlesinger F, Drenkow J, Zaleski C, Jha S, Batut P, Chaisson M, Gingeras TR. 2013. STAR: ultrafast universal RNA-seq aligner. Bioinformatics 29: 15–21.

Frankish A, Diekhans M, Ferreira AM, Johnson R, Jungreis I, Loveland J, Mudge JM, Sisu C, Wright J, Armstrong J et al. 2019. GENCODE reference annotation for the human and mouse genomes. Nucleic Acids Res 47: D766–D773.

Kozak M. 1987. An analysis of 5’-noncoding sequences from 699 vertebrate messenger RNAs. Nucleic Acids Res 15: 8125–8148.

Kuksa PP, Amlie-Wolf A, Katanic Z, Valladares O, Wang LS, Leung YY. 2019. DASHR 2.0: integrated database of human small non-coding RNA genes and mature products. Bioinformatics 35: 1033–1039.

Kurmangaliyev YZ, Sutormin RA, Naumenko SA, Bazykin GA, Gelfand MS. 2013. Functional implications of splicing polymorphisms in the human genome. Hum Mol Genet 22: 3449–3459.

Landrum MJ, Lee JM, Benson M, Brown GR, Chao C, Chitipiralla S, Gu B, Hart J, Hoffman D, Jang W et al. 2018. ClinVar: improving access to variant interpretations and supporting evidence. Nucleic Acids Res 46: D1062–D1067.

Lee CM, Barber GP, Casper J, Clawson H, Diekhans M, Gonzalez JN, Hinrichs AS, Lee BT, Nassar LR, Powell CC et al. 2020. UCSC Genome Browser enters 20th year. Nucleic Acids Res 48: D756–D761.

O’Leary NA, Wright MW, Brister JR, Ciufo S, Haddad D, McVeigh R, Rajput B, Robbertse B, Smith-White B, Ako-Adjei D et al. 2016. Reference sequence (RefSeq) database at NCBI: current status, taxonomic expansion, and functional annotation. Nucleic Acids Res 44: D733–745.

Pal LR, Moult J. 2015. Genetic Basis of Common Human Disease: Insight into the Role of Missense SNPs from Genome-Wide Association Studies. J Mol Biol 427: 2271–2289.

Quast C, Pruesse E, Yilmaz P, Gerken J, Schweer T, Yarza P, Peplies J, Glockner FO. 2013. The SILVA ribosomal RNA gene database project: improved data processing and web-based tools. Nucleic Acids Res 41: D590–596.

Schneider VA, Graves-Lindsay T, Howe K, Bouk N, Chen HC, Kitts PA, Murphy TD, Pruitt KD, Thibaud-Nissen F, Albracht D et al. 2017. Evaluation of GRCh38 and de novo haploid genome assemblies demonstrates the enduring quality of the reference assembly. Genome Res 27: 849–864.

Semler O, Garbes L, Keupp K, Swan D, Zimmermann K, Becker J, Iden S, Wirth B, Eysel P, Koerber F et al. 2012. A mutation in the 5’-UTR of IFITM5 creates an in-frame start codon and causes autosomal-dominant osteogenesis imperfecta type V with hyperplastic callus. Am J Hum Genet 91: 349–357.

Shibata N, Ohoka N, Sugaki Y, Onodera C, Inoue M, Sakuraba Y, Takakura D, Hashii N, Kawasaki N, Gondo Y et al. 2015. Degradation of Stop Codon Read-through Mutant Proteins via the Ubiquitin-Proteasome System Causes Hereditary Disorders. J Biol Chem 290: 28428–28437.

Starck SR, Jiang V, Pavon-Eternod M, Prasad S, McCarthy B, Pan T, Shastri N. 2012. Leucine-tRNA initiates at CUG start codons for protein synthesis and presentation by MHC class I. Science 336: 1719–1723.

Varki A, Altheide TK. 2005. Comparing the human and chimpanzee genomes: searching for needles in a haystack. Genome Res 15: 1746–1758.

von Bohlen AE, Bohm J, Pop R, Johnson DS, Tolmie J, Stucker R, Morris-Rosendahl D, Scherer G. 2017. A mutation creating an upstream initiation codon in the SOX9 5’ UTR causes acampomelic campomelic dysplasia. Mol Genet Genomic Med 5: 261–268.

Wang SH, Hsiao CJ, Khan Z, Pritchard JK. 2018. Post-translational buffering leads to convergent protein expression levels between primates. Genome Biol 19: 83.

Yang Z. 2007. PAML 4: phylogenetic analysis by maximum likelihood. Mol Biol Evol 24: 1586–1591.

Yngvadottir B, Xue Y, Searle S, Hunt S, Delgado M, Morrison J, Whittaker P, Deloukas P, Tyler-Smith C. 2009. A genome-wide survey of the prevalence and evolutionary forces acting on human nonsense SNPs. Am J Hum Genet 84: 224–234.

